# Alpha rhythms reveal when, where and how memories are retrieved

**DOI:** 10.1101/708602

**Authors:** María Carmen Martín-Buro, Maria Wimber, Richard N. Henson, Bernhard P. Staresina

**Author notes:** **Corresponding and Lead author:** Bernhard P. Staresina.

## Abstract

Our memories for past experiences can range from vague recognition to full-blown recall of associated details. Neuroimaging research has tried to understand the brain mechanisms underlying qualitatively different memories for decades (Yonelinas, 2002). On the one hand, Electroencephalography (EEG) has shown that recall signals unfold a few hundred milliseconds after simple recognition and are hallmarked by sustained voltage deflections over left posterior sensors (Herron, 2007; Johansson & Mecklinger, 2003; Mecklinger, Rosburg, & Johansson, 2016; Rugg & Curran, 2007). However, sensor-based analyses only provide limited insights into the supporting brain networks. On the other hand, functional magnetic resonance imaging (fMRI) has revealed a ‘core recollection network’ centred on posterior parietal and medial temporal lobe (MTL) regions (Hayama, Vilberg, & Rugg, 2012; Johnson, Suzuki, & Rugg, 2013; King, de Chastelaine, Elward, Wang, & Rugg, 2015; Rugg, Johnson, & Uncapher, 2015; Rugg & Vilberg, 2013; Thakral, Benoit, & Schacter, 2017). However, due to the relatively poor time resolution of fMRI, the temporal dynamics of these regions during retrieval remain largely unknown. In order to overcome these modality-specific limitations, we here used Magnetoencephalography (MEG) in a verbal episodic memory paradigm assessing correct rejection (CR) of lures, item recognition (IR) of old words and associative recall (AR) of paired target words. We found that power decreases in the alpha frequency band (10-12 Hz) systematically track different mnemonic outcomes in both time and space: Over left posterior sensors, alpha power decreased in a stepwise fashion from 500 ms onward, first from CR to IR and then from IR to AR. When projecting alpha power into source space, the ‘core recollection network’ known from fMRI studies emerged, including posterior parietal cortex (PPC) and hippocampus. While PPC showed a linear change across conditions, hippocampal effects were specific to recall. Critically, the hippocampal recall effect emerged ∼200 ms before the PPC recall effect, suggesting a bottom-up recall signal from hippocampus to PPC. Our data thus link engagement of the core recollection network to the temporal dynamics of episodic memory and suggest that alpha rhythms constitute a fundamental oscillatory mechanism revealing when, where and how our memories are retrieved.

**Highlights:**

- Alpha rhythms distinguish between different retrieval outcomes
- Alpha power time courses track item recognition and associative recall
- Source alpha power decreases track the fMRI core recollection network
- Hippocampal recall signal precedes parietal signal

## Results and Discussion

Episodic memory, our ability to remember past events and experiences, is a key pillar of cognition and behaviour. Intriguingly though, some memories remain faint, eliciting a sense of familiarity at best, while others are vivid and bring back a wealth of associations (Yonelinas, 2002). Investigation of the neural mechanisms supporting memory recall has been ignited by EEG studies revealing a left posterior ‘old/new’ effect, i.e., a difference in slow event-related potentials (ERPs) over left posterior sensors unfolding between 500 and 1000 ms after cue onset (Sanquist et al., 1980; for a review see Rugg and Curran, 2007). In parallel, fMRI studies have consistently shown a core brain network, featuring parietal and medial temporal regions, differentially engaged during successful recollection (Hayama et al., 2012; Rugg & Vilberg, 2013). However, due to inherent limitations of both methods (relatively poor spatial resolution of scalp ERPs, poor temporal resolution of fMRI), it is unclear whether the cue-evoked ERPs reflect engagement of the core recollection network and whether engagement of the fMRI network is temporally linked to the moment of retrieval, as opposed to pre-stimulus/preparatory deployment of attention or post-retrieval monitoring (Levy, 2012; Sestieri, Shulman, & Corbetta, 2017). Moreover, it is challenging to disentangle the temporal dynamics within the recollection network with fMRI, allowing only speculation about whether parietal regions drive the hippocampus in a top-down manner during successful recall or whether the hippocampus provides a bottom-up signal to parietal regions (Ciaramelli, Grady, & Moscovitch, 2008; Vilberg & Rugg, 2008; Wagner, Shannon, Kahn, & Buckner, 2005). Direct intracranial recordings would provide the desired temporal and spatial resolution, but comprehensive coverage of both parietal and mediotemporal areas is rare and advanced retrieval paradigms (probing different types of memory) are challenging to conduct with patients (Foster, Rangarajan, Shirer, & Parvizi, 2015; Gonzalez et al., 2015).

That said, one viable means of integrating the strengths of EEG and fMRI recordings might be the examination of oscillatory patterns in the alpha frequency band (8-12 Hz). On the one hand, simultaneous EEG-fMRI recordings have revealed a strong link between blood-oxygenation-level-dependent (BOLD) signal increases and decreases in alpha power (‘desynchronisation’) (Laufs et al., 2003; Meltzer, Negishi, Mayes, & Constable, 2007; Moosmann et al., 2003; Scheeringa et al., 2011). On the other hand, modelling and empirical work suggests that slow and late ERPs might reflect asymmetric amplitude fluctuations in the alpha band, such that e.g. oscillatory peaks become more pronounced than troughs over time (Mazaheri & Jensen, 2008). We thus hypothesised that alpha desynchronization not only differentiates between different types of episodic retrieval in the time domain (from ∼500 ms onward), but that this effect spatially maps onto the core recollection network, thus pinpointing its purported role in peri-stimulus retrieval.

Capitalising on the increased spatial resolution of MEG over EEG (Baillet, 2017; Lopes da Silva, 2013), we employed a memory retrieval paradigm (Figure 1) in which participants (n=15) indicated whether a given word was (i) new, (ii) old but they could not recall the paired associate, or (iii) old and they also recalled the paired associate. In the latter case, a second screen appeared in which participants indicated which of three first-and-last-letter combinations corresponded to the target paired associate.

**Figure 1.**
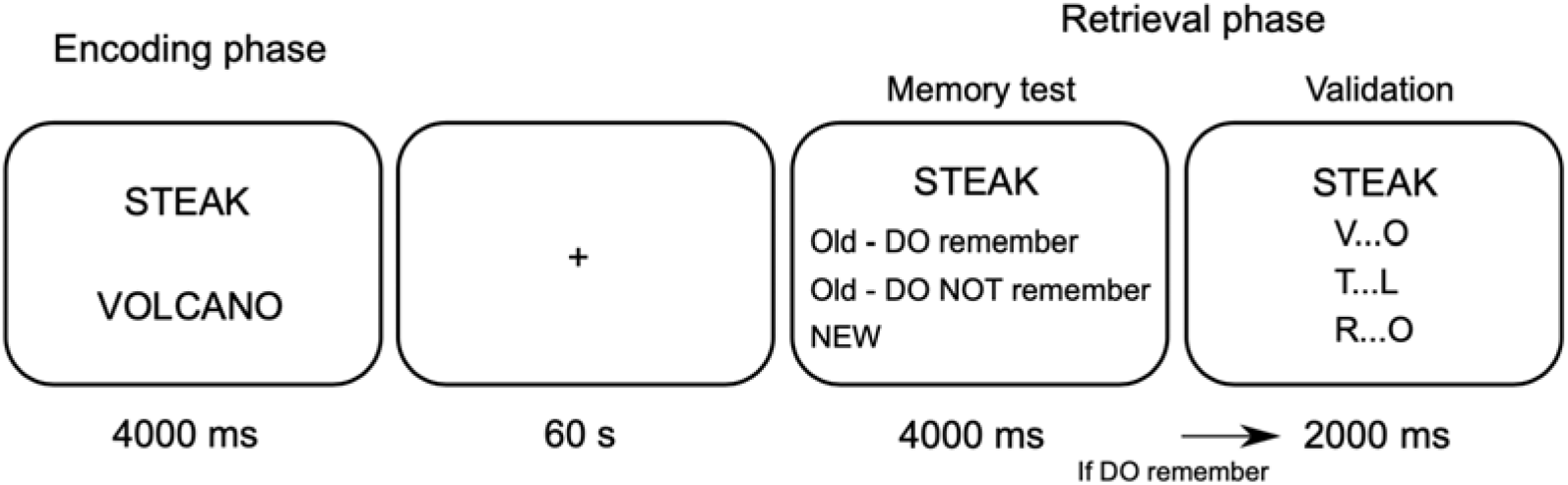
Experimental procedure. During the study phase (‘encoding’), participants saw word pairs under deep or shallow processing tasks. During the subsequent test phase (‘retrieval’), one word of the previously presented pairs was shown, intermixed with unstudied new words (‘lures’). Participants indicated with one button press whether they thought the given word was new, the word was old but they did not remember the paired associate or the word was old and they recalled the paired associate. In the latter case, a second screen appeared to validate recall accuracy, providing three first-last letter combinations of which one corresponded to the target association. Analyses focused on correct identification of lures (correct rejection, CR), correct identification of old words without recalling the paired associate (item recognition; IR) and correct identification of old words along with correctly recalling the paired associate (associative recall; AR).

Focusing on correct memory outcomes, our three conditions of interest were (i) correct rejection of new words (CR), (ii) correct identification of old words, without recalling the paired associate (item recognition memory, IR) and (iii) correct identification of old words along with correct recall of the paired associate (associative recall, AR). In terms of nomenclature, we define an item recognition effect as the difference between IR and CR and an associative recall effect as the difference between AR and IR. We used a levels-of-processing manipulation during encoding (Craik & Lockhart, 1972) to yield balanced numbers of both IR and AR alongside CR trials (see STAR Methods and Table 1). The overall rate of HITs (collapsing IR and AR) minus false alarms was .59, indicating high levels of recognition memory. The proportion of correct forced choices during the validation task was .96 (SEM = .01), indicating high levels of paired associate recall after the initial AR response. Reaction times (RTs) differed significantly across our conditions of interest: RTs for Hits were significantly longer than for CR (t_14_ = 3.26; p = .005), and for IR compared to AR (t_14_ = 3.87; p = .001).

**Table 1.** Retrieval accuracy and reaction times. For Correct rejections and Hits, proportion denotes proportion of all new (112) and old (224) trials.

|  | Proportion |  | Reaction times (s) |  |
| --- | --- | --- | --- | --- |
|  | Mean | SEM | Mean | SEM |
| Correct rejections | 0.87 | 0.03 | 1.49 | 0.06 |
| Hits | 0.72 | 0.04 | 1.76 | 0.07 |
| Associative recall | 0.45 | 0.05 | 1.53 | 0.07 |
| Item recognition | 0.55 | 0.05 | 1.92 | 0.09 |

### Alpha rhythms track time courses of item recognition and associative recall

Given the RT distribution across trial types (Table 1), we restricted our sensor space analysis to the first 2 seconds after cue onsets (longest average RT of 1.92 s). To identify - in one step - time points, frequencies and sensors modulated by memory outcome, we first conducted a repeated-measures ANOVA with the factor Memory (CR, IR, AR) on time-frequency representations (TFRs, relative power change) across sensors (planar gradiometers). Results showed a significant effect surviving cluster-based correction for multiple comparisons (Maris & Oostenveld, 2007) (p < .001). As shown in Figure 2A, the effect was centred at left posterior sites, spanning a time window of 0.7-2 s and a frequency range from 8-20 Hz, with a distinctive peak from 10-12 Hz (alpha frequency range). To maximise sensitivity, subsequent analyses focus on this 10-12 Hz band, but results remain stable when including a wider range of frequencies and sensor selections (Figure S1). Likewise, our first time course analyses (Figure 2C and 2D) focus on the 10 contiguous left posterior sensors with the maximal F values to link results to the previous M/EEG literature, but subsequent source space analyses capitalise on the entire sensor array. First, extracting the corresponding power values for the three memory conditions, post hoc pairwise tests revealed a stepwise decrease in alpha power from CR to IR (t_14_ = −8.41, p < .001) and from IR to AR (t_14_ = −4.75, p < .001) (Figure 2B). These results extend previous findings of left posterior alpha power distinguishing between correctly recognised old and new items (Hanslmayr, Staudigl, & Fellner, 2012), now showing that it further distinguishes between item recognition and associative recall.

**Figure 2.**
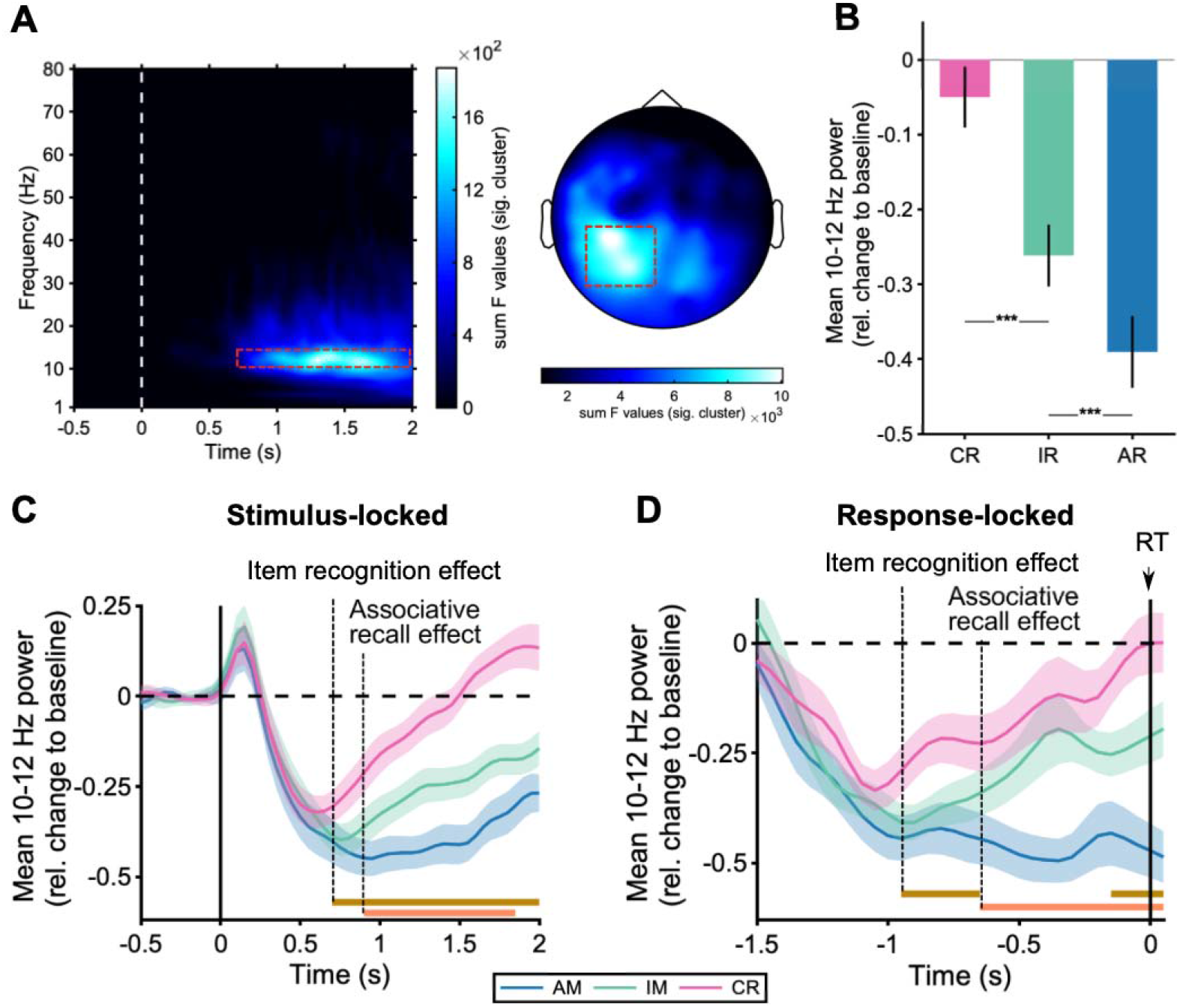
Sensor space results. (A) ANOVA results for the comparison of CR, IR and AR TFRs revealed a significant cluster from 0.7-2 s at left posterior sensors with a peak at 10-12 Hz. TFR plot (*left*) depicts the sum of F-values across all significant sensors of the cluster. Topoplot (*right*) shows the sum of F-values across all significant time/frequency bins of the cluster. (B) Mean (+/−SEM) alpha power for each memory condition collapsed across left posterior sensors from 0.7-2 s in the 10-12 Hz frequency range (red dashed boxes in A), showing a relative power decrease (‘desynchronization’) modulated by memory outcome. ***: p < .001, paired samples t test. (C) Alpha power (10-12 Hz) time courses, collapsed across left posterior sensors (cf. Figure 2A). (C) Stimulus-locked and (D) Response-locked averages across participants (+/−SEM). Dashed vertical lines highlight onsets at which item recognition memory effects (IR vs. CR) and associative recall effects (AR vs. IR) effects unfold, and brown and orange horizontal lines depict the significant clusters for the respective paired-samples T-tests (all p < .005)

Do IR and AR effects in the alpha band unfold at different latencies, tracking the delay of recollection relative to familiarity-based recognition (Yonelinas, 2002) or the gradual accumulation of mnemonic evidence (Wagner et al., 2005), respectively? To address this question, we examined the time courses of alpha power at left posterior sensors for CR, IR and AR. As shown in Figure 2C, an IR effect emerged at 700 ms post cue onset. Next, with a delay of ∼150 ms, an AR effect emerged as a significant decrease in alpha power for AR relative to IR. To ensure that our effects do not reflect post-retrieval processes (e.g., idling or monitoring, *see below), we* repeated the timecourse analysis with response-locked rather than stimulus-locked data, thereby accounting for different response latencies across memory conditions (Table 1). Indeed, results confirmed that the differential IR and AR effects unfolded well before the behavioural response: The IR effect emerged ∼950 ms prior to the response, followed by an AR effect onsetting ∼650 ms prior to the response (Figure 2D). Finally, to quantify whether alpha power decreases peaked at different latencies for CR, IR and AR, we derived participant-specific time points of maximal alpha power decrease at sensors highligted in Figure 2A (dashed red square). Mean peak latencies were 677 ms (SEM = 62 ms) for CR, 877 ms (SEM = 95 ms) for IR and 1113 ms (SEM = 108 ms) for AR. A repeated measures ANOVA with the factor Memory (CR, IR, AR) on these peaks confirmed a significant main effect (F_(2,28)_ = 7.44, p = .002) with a significant linear term (F_(1,14)_ = 11.31, p = .005). Post hoc pairwise tests revealed a trending effect for CR vs. IR (t_14_ = −1.81, p = .09) and a significant difference for IR vs. AR (t_14_ = - 2.5, p = .02).

### Alpha rhythms track engagement of the core recollection network

As shown in Figure 2A, the sensor-level alpha effects were most pronounced over left posterior sites. While this topography is well in line with a host of ERP studies revealing a left posterior recognition memory effect (Sanquist et al., 1980; for a review see Rugg and Curran, 2007), more recent fMRI investigations of recognition memory have consistently revealed a ‘core-recollection’ network, including posterior parietal cortex (PPC) and medial temporal lobe regions. We next projected our data into source space and first focused our source level analysis on the 0.7 to 2 s post-stimulus time window and 10-12 Hz frequency band to best capture the memory effects previously found in the sensor-space analysis (Figure 2) (see STAR Methods). Thresholding the statistical F map from an omnibus ANOVA at p < .05 (corrected) revealed prominent peaks in medial and lateral PPC (including precuneus, retrosplenial cortex, superior and inferior parietal lobule), lateral temporal cortex (LTC), as well as the hippocampus (Figure 3A and Figure S2).

**Figure 3.**
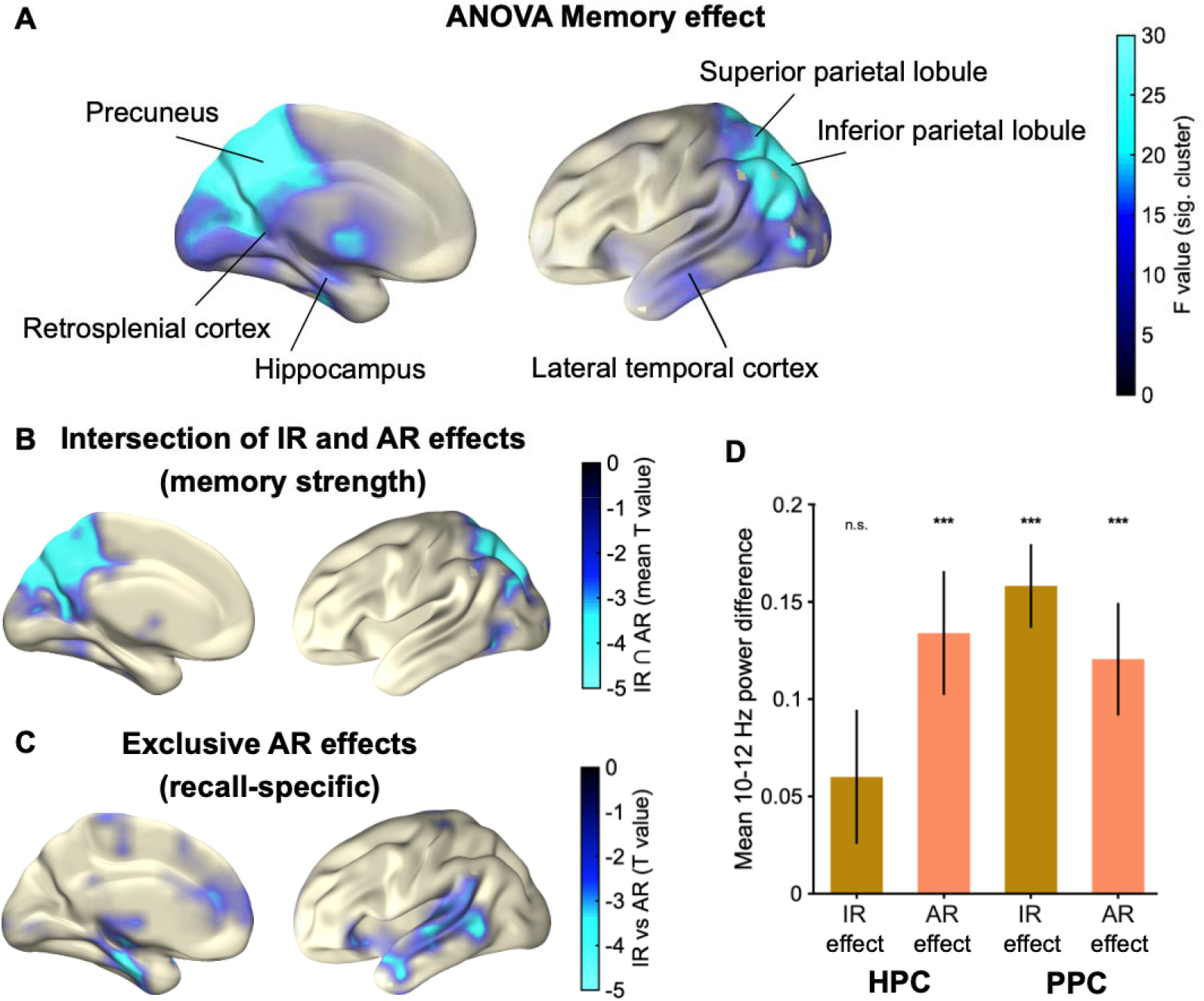
Source reconstruction. (A) Significant cluster resulting from the ANOVA in the 10-12 Hz alpha band from 0.7 to 2 s. (B) Regions scaling with memory strength (CR < IR < AR), revealed via inclusive masking of condition comparisons (intersection of IR vs. CR and AR vs. IR) in the 10-12 Hz alpha band and from 1 to 1.5 s. Colorbar indicates the mean T values across the IR effect (CR < IR) and the AR effect (IR < AR). (C) Exclusive AR effects (recall-specific) map (1 to 1.5 s) indicates areas showing an AR effect (AR > IR, p < .05, corrected) and no IR effect (IR > CR, p < .1, uncorrected). Colorbar indicates the T value for the AR effect. Labelling of brain regions is based on the Automated Anatomical Labelling (AAL) atlas (Tzourio-Mazoyer et al., 2002). (D) Post-hoc comparisons for independently defined hippocampus (HPC) and PPC regions of interest (AAL atlas). Mean (+/−SEM) IR effect (CR vs IR) and AR effect (IR vs AR) from 1-1.5 s in the 10-12 Hz frequency range. ***: p < .001, one-sample t test of effects vs. 0.

Within the core recollection network, fMRI studies have consistently revealed functional dissociations, such that PPC regions track memory strength in a linear fashion (here: CR < IR < AR), whereas the hippocampus selectively supports recall-based memory (CR = IR < AR) (Hayama et al., 2012; Vilberg & Rugg, 2014). To test whether alpha power source localisation is able to track these qualitative differences, we applied in- and exclusive masking analyses on the source reconstructed data from 1-1.5 s, where both the IR effect and the AR effect were observable in sensor space (Figure 2C), thus ensuring a fair comparison between conditions. First, to reveal regions that show a stepwise increase in alpha power desynchronization, we inclusively masked the IR effect (IR > CR) with the AR effect (AR > IR), with both effects thresholded at P < .05 (corrected). The conjoint effect revealed medial and lateral PPC (Figure 3B). Next, to highlight regions specifically supporting recall in our paradigm, we conducted the contrast of AR > IR (P < .05, corrected) and excluded regions that would also show an IR effect (IR > CR), liberally thresholded at P < .1, uncorrected. Note that the more liberal the exclusive mask (IR > CR), the more conservative the specificity to the initial contrast (AR > IR). Strikingly, this procedure revealed the hippocampus along with lateral temporal cortex and prefrontal cortex (Figure 3C). To complement the masking approach, we additionally extracted alpha power from hippocampus and PPC in anatomically defined regions of interest (based on the AAL atlas). A Region (Hippocampus, PPC) x Memory (CR, IR, AR) repeated-measures ANOVA on 10-12 Hz alpha power from 1-1.5s showed a significant main effect of Region (F_(1,14)_= 17.07, p = .001), a significant main effect of Memory (F_(1,14)_ = 30.45, p < .001) and, critically, a significant Region x Memory interaction (F_(1,14)_= 5.72, p = .017). Follow-up comparisons of IR and AR effects (Figure 3D) confirmed that PPC showed both an IR and an AR effect (both t_(14)_ > 4.20, p < .001), whereas the hippocampus showed an AR effect (t_(14)_= 4.10, p < .001) but no IR effect (t_(14)_,= 1.17, p = .10). In fact, the IR effect was significantly greater in PPC than in the hippocampus (t_(14)_ = 3.31, p = .005). In sum, our source reconstruction analyses revealed a remarkable overlap between the fMRI core recollection network and the regional pattern of alpha power decreases. Not only does this elucidate alpha desynchronization as an oscillatory mechanism governing the contribution of different brain regions to different types of memory retrieval, but it also opens the window for examining the temporal dynamics within the recollection network.

### Alpha rhythms reveal different temporal profiles within the core recollection network

Recent fMRI studies have begun to shed some light on the temporal profiles of PPC and hippocampal engagement during retrieval. By varying the interval of maintaining a recalled episodic detail, Vilberg and Rugg (2014) were able to show that hippocampal engagement during successful recall was transient, whereas PPC engagement was sustained and covaried in time with the maintenance interval (see Thakral, Benoit, et al., 2017 for similar results in an episodic future simulation paradigm). While this pattern is consistent with the scenario that PPC mechanisms are deployed to work with mnemonic content provided by the hippocampus, temporal precedence of a hippocampal relative to a PPC recall effect would provide more stringent evidence for this notion. We thus extracted the alpha power time course from PPC and hippocampus in order to examine a possible temporal dissociation in the onset of these regions’ AR effect (see STAR Methods). Results indicate that the hippocampal AR effect started at 700 ms after stimulus onset, whereas the PPC AR effect set in with a 200ms delay, at 900 ms after stimulus onset. These patterns point to a recall-specific signal in the hippocampus, which is followed by PPC recruitment, with the latter possibly reflecting the additional amount of memory strength/mnemonic detail (Wagner et al., 2005; Rugg & Vilberg, 2013) and/or attention to memory (Ciaramelli, Grady, Levine, Ween, & Moscovitch, 2010). It deserves explicit mention though that latency differences of significant effects are not equivalent to significant differences of effect latencies. That is, although the hippocampal AR effect is statistically significant at an earlier time point than the PPC AR effect, this does not mean that the observed latency difference of the effect across regions (here, 200 ms) is statistically significant.

### Alpha desynchronization indicates when, where and how memories are retrieved

Our results show that alpha power desynchronization unifies the temporal and spatial profiles of recall via a single physiological mechanism. Importantly, this allowed us to reveal that a hippocampal recall signal precedes the PPC recall signal, pointing to a role of PPC in representing/manipulating mnemonic content provided by the hippocampus (see below).

Despite the long history of M/EEG studies on recognition memory (Sanquist et al., 1980; for reviews, see Mecklinger, 2000; Rugg and Curran, 2007), only a few have examined oscillatory patterns related to different memory outcomes (Burgess & Gruzelier, 2000; Khader & Rösler, 2011; Michelmann, Bowman, & Hanslmayr, 2016; Vogelsang, Gruber, Bergström, Ranganath, & Simons, 2018; Waldhauser, Braun, & Hanslmayr, 2016), albeit without explicitly distinguishing associative recall from item recognition. Our current paradigm allowed us to directly probe the oscillatory mechanisms that support these different memory signals (Figure 1). As shown in Figure 2, results revealed that left posterior alpha desynchronization not only tracked simple old/new recognition memory (IR vs. CR), but further distinguished between old/new recognition and associative recall (AR vs. IR). Indeed, time course analyses (Figure 2C and D) confirmed the temporal offset between an earlier IR effect (starting at ∼700 ms after cue onset) followed by a later AR effect (starting at ∼900 ms after cue onset) (Rugg & Yonelinas, 2003; Yonelinas, 2002). We note that the onset latency of the IR effect is markedly later than the FN400 component (negative signal deflection over frontal sites around 400 ms) traditionally linked to familiarity-based recognition (Curran, 2000; Düzel, Yonelinas, Mangun, Heinze, & Tulving, 1997; Johansson & Mecklinger, 2003; Rugg & Curran, 2007). Given the spatial and temporal extent of the left posterior alpha effect (Figure 2A), less extended/more local effects might have been overshadowed by the cluster-based correction method. We thus conducted a more targeted analysis, directly contrasting IR vs. CR alpha power in source space from 300-500 ms. Indeed, as shown in Figure S3, this revealed a significant familiarity effect in prefrontal and anterior MTL cortical regions. We thus suggest that the stepwise change in alpha power at left posterior sites, including a stepwise delay in peak latencies (CR < IR < AR), reflects the gradual accumulation of memory strength/mnemonic evidence (Wagner et al., 2005). In any case, considering the potential link between amplitude fluctuations in the alpha band and sustained ERP deflections (Mazaheri & Jensen, 2008), our data raise the possibility that at least some of the classic ERP recognition effects reflect condition-specific differences in alpha power.

In a separate line of research, fMRI studies on recognition memory have consistently shown engagement of a particular set of brain regions in recall-based memory, including lateral/medial parietal and temporal regions. The robustness of these regions’ engagement across numerous paradigms has given rise to the notion that they represent a core recollection network (Rugg & Vilberg, 2013). However, given the relatively poor temporal resolution of fMRI, it has been challenging to pinpoint the exact cognitive (sub)processes during recognition these regions support. Accordingly, while some accounts posit that this network represents informational content during retrieval (Johnson et al., 2013; Rugg & Vilberg, 2013; Thakral et al., 2017; Vilberg & Rugg, 2014), others highlight – particularly regarding parietal contributions – pre-retrieval (Cabeza, 2008), peri-retrieval (Haramati, Soroker, Dudai, & Levy, 2008; Shimamura, 2011; Wagner et al., 2005) or post-retrieval (Ciaramelli et al., 2010) operations (for reviews see Levy, 2012; Sestieri et al., 2017). Projecting our sensor data into source space, we found a striking overlap of our alpha power memory effects with the core recollection network (Figure 3). The pair-wise comparisons showed that both IR and AR effects map onto bilateral (superior/inferior parietal lobule) and medial parietal cortex (precuneus/retrosplenial cortex; see also Bergström, Henson, Taylor, & Simons, 2013). Conversely, hippocampus and medial prefrontal cortex showed specific engagement for AR (Figure 3C). The topographical correspondence of our alpha power decreases with BOLD increases commonly found in fMRI studies on recognition memory adds to a number of EEG-fMRI studies showing a tight coupling of these two measures (Laufs et al., 2006) and suggests that alpha power can, at least in some cases, be used as a time-resolved proxy for BOLD activation.

The yoking of alpha desynchronization effects with the fMRI recollection network opens insights into this network’s temporal profile and informs theories on hippocampal and PPC contributions to memory retrieval. First, taking sensor space (Figure 2C) and source space temporal dynamics (Figure 4) together, the memory effects clearly emerged after cue onset but well before the mnemonic decision (mean RT = 1.76), pointing to peri-retrieval engagement of the recollection network rather than pre-stimulus preparatory or post-retrieval monitoring/decision making functions. Moreover, across hippocampus and PPC, the source power time courses (Figure 4) suggest that recall success is initiated by the hippocampus and PPC might govern the ensuing accumulation of mnemonic evidence and/or provide an ‘episodic buffer’ (Baddeley, 2000; Hayama et al., 2012; Rugg & Vilberg, 2013; Shimamura, 2011).

**Figure 4.**
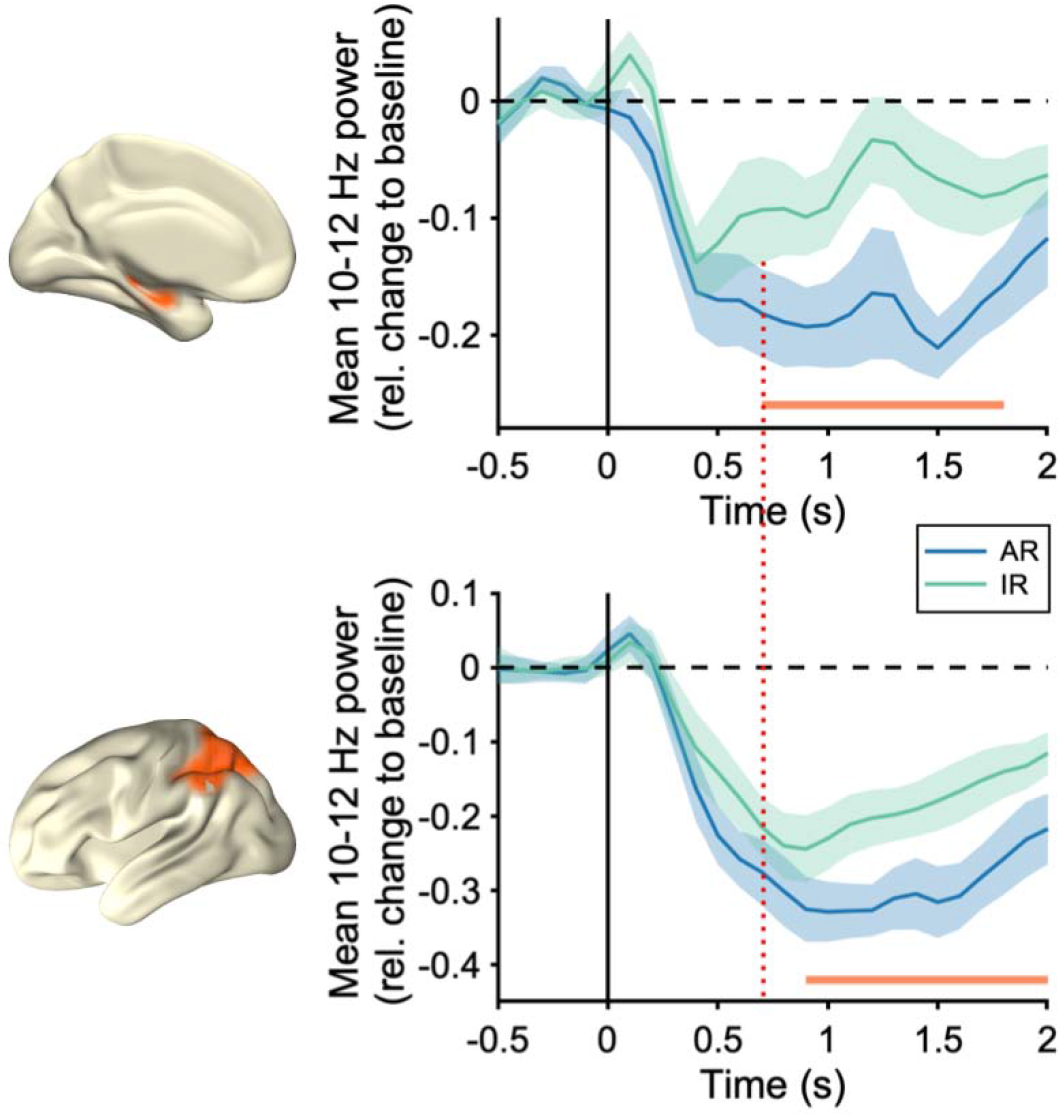
Hippocampus and PPC alpha source power time courses. 10-12 Hz alpha source power for AR and IR conditions. Brain maps depict the regions of interest selected for this analysis based on the Automated Anatomical Labelling (AAL) atlas (Tzourio-Mazoyer et al., 2002). *Top* panel depicts alpha power time courses in the hippocampus. *Bottom* panel includes inferior and superior parietal lobules. Orange horizontal lines represent the significant time points of the IR vs AR t-test (cluster corrected).

The link between parietal alpha power decreases and the accumulation of mnemonic evidence also aligns with a recent model suggesting that low frequency desynchronization reflects the amount of information represented by a given region (Hanslmayr et al., 2012; Hanslmayr et al., 2016) rather than mere activation (Pfurtscheller & Lopes, 1999) or disinhibition (Jensen & Mazaheri, 2010; Klimesch, 1996). In the hippocampus, the alpha power decrease for AR vs. IR may again reflect an increase in information, though it deserves explicit mention that no differences were observed between IR and CR (Figure 4B and C), consistent with a selective role of the hippocampus in associative/relational retrieval operations (Davachi, 2006; Eichenbaum, Yonelinas, & Ranganath, 2007; Mayes, Montaldi, & Migo, 2007). The MEG alpha power decrease in the hippocampus observed here is remarkably similar to that shown in a recent iEEG study using direct hippocampal recordings (Staresina et al., 2016), both in the frequency range and effect latency. In that study, the alpha power decreases for associative vs. non-associative retrieval (similar to AR vs. IR here) was preceded by a gamma power (∼50-90 Hz) increase at 500 ms. One plausible scenario might be that the gamma power increase at 500 ms reflects hippocampal pattern completion processes, with the ensuing alpha power decrease reflecting an increase in information emerging from this process. Of course caution is warranted when interpreting MEG effects in deep anatomical sources such as the hippocampus, but besides the convergence of results with direct iEEG recordings, our findings add to a growing body of evidence of discernible MEG effects in the hippocampus (for review, see Pu et al., 2018).

Finally, while our effects were most prominent in the alpha frequency band (Figure 2A), it is important to note that other low frequency bands, particularly theta (4-8 Hz), have also been linked to memory processes. For instance, Osipova et al. (Osipova et al., 2006) found theta increases for HITs relative to CRs in an image recognition paradigm. Interestingly, though, this effect was localised to occipital cortex and already started 300 ms post cue onset. Theta power increases have also been linked to hippocampal retrieval process in iEEG recordings (Burke et al., 2014), though that study employed a free recall rather than a recognition memory/cued recall paradigm. Another recent study combined MEG recordings with continuous theta burst stimulation (cTBS) during an autobiographical memory task (Hebscher, Meltzer, & Gilboa, 2019) and found theta power increases and theta-gamma coupling in the core recollection network. Together, this raises the possibility that different functional networks, recruited by different memory demands, are grouped by different frequency bands, and an important challenge for future studies will thus be to delineate the roles of theta power increases vs. alpha power decreases in service of episodic retrieval (Hanslmayr et al., 2016).

To conclude, our understanding of recognition memory thus far relied upon separate lines of research capitalizing on either temporal or spatial signal properties. Our study now suggests that alpha rhythms represent a single oscillatory mechanism tracking where and when associative memory unfolds in space and time, linking differential engagement of the hippocampus and parietal cortex to episodic retrieval processes with millisecond precision.

## STAR methods text

### KEY RESOURCES TABLE

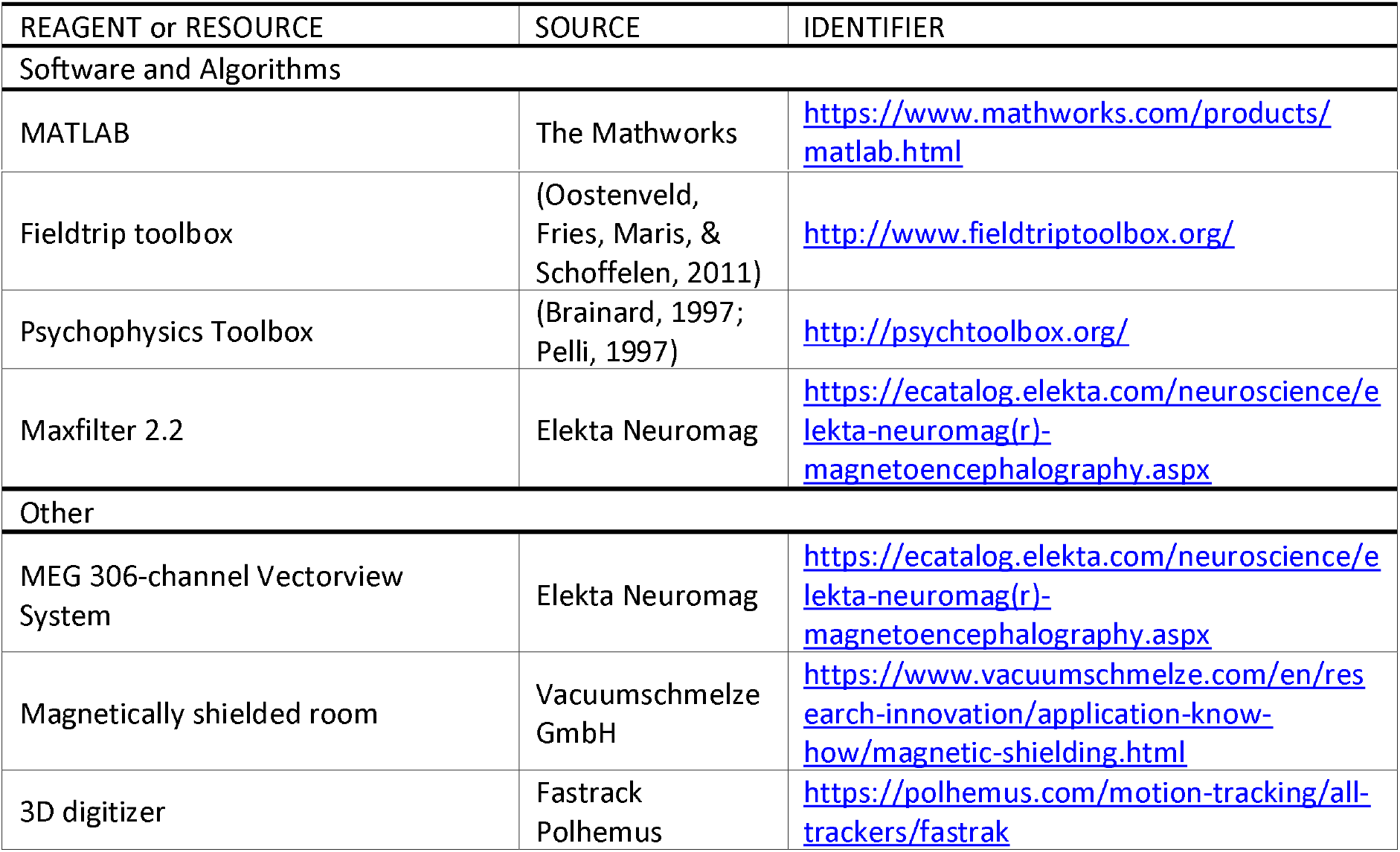

### CONTACT FOR REAGENT AND RESOURCES SHARING

Further information and request for resources and reagents should be directed to and will be fulfilled by the Lead Author Bernhard P. Staresina

### DATA AND SOFTWARE AVAILABILITY

For access to data and software, please contact Bernhard P. Staresina

### EXPERIMENTAL MODEL AND SUBJECT DETAILS

We designed a repeated-measures experiment with a sample size of 15 healthy right-handed participants (9 females; mean age: 24 years, range: 18-37) who gave written informed consent. All procedures were approved by University of Cambridge Psychology Research Ethics Committee.

### METHODOLOGY DETAILS

#### Behavioural analysis

The experiment was programmed in MATLAB using the Psychophysics Toolbox extensions (Brainard, 1997; Pelli, 1997). Behavioural data were processed and analysed using custom-written MATLAB code.

#### Experimental paradigm

The experiment was conducted inside the MEG shielded room with the participant seated upright. A schematic diagram of the experimental paradigm is shown in Figure 1. Participants completed eight encoding-retrieval runs with 60 seconds before and after each encoding phase in which they were asked to look at a central fixation cross. During encoding, participants were presented with pairs of English nouns. In order to obtain experimental leverage on different memory outcomes (item and associative memory), we used two different encoding tasks that varied in the depth of processing: a *syllable task* in which participants indicated how many of the two words contained 2 syllables (0, 1 or 2; *shallow encoding*), and an *imagery task* in which participants vividly imagined the two objects interact and indicated their imagery success (low, medium, high; *deep encoding*). Each word pair remained on the screen for 4 seconds regardless of the participant’s response. Incidental to the encoding task, a flickering background, flickering at 8.6 or 12 Hz, was presented on the left or right side of the screen which participants were instructed not to pay attention to. The flicker manipulation during encoding is beyond the scope of the current manuscript, but counterbalancing ensured that deep and shallow encoding trials were equally often presented with both flicker rates and at both visual hemifields. Each encoding block contained 28 word pairs, with deep and shallow tasks alternating every 7 trials. During the subsequent retrieval block, participants were presented with one randomly chosen word from each of the 28 previously seen pairs as well as 14 novel nouns. First, participants indicated if the word was (i) old and they also remembered the paired associate, (ii) old but they could not remember the paired associate, or (iii) new. The response was collected with a single button press and the word remained in the screen during 4 seconds regardless of the participant’s response. When the first option was selected, a validation screen of 2 seconds of duration appeared and the participants had to choose which of three first-and-last letter combinations corresponded to the remembered paired associate. This two-step structure served as a means of objective validation while holding the stimulus display and response options constant for the initial 4 seconds of the trial. Preceding each trial, a fixation cross was displayed during a jittered intertrial interval of 850 to 1150ms. For subsequent analyses, the following three conditions of interest were defined: Associative Recall (AR; trials in which participants indicated they recognized an old word and recalled the paired associate, followed by a correct response during verification); Item Recognition (IR; trials in which participants indicated they recognized an old word but did not recall the paired associate) and Correct Rejection (CR; trials in which participants correctly identified new items). In order to restrict our analyses to correct memory trials, we excluded Misses (trials in which old items were incorrectly identified as new), False Alarms (trials in which new items were incorrectly identified as old) and trials in which participants first indicated they recalled the word plus its paired associate but then gave an incorrect response during verification.

#### MEG Recordings

Data were recorded in a magnetically shielded room using a 306-channel VectorView MEG system (Elekta Neuromag, Helsinki). Data were sampled at 1 kHz with a highpass filter of 0.03 Hz. Only the 204 planar gradiometers were used in the analysis. Head position inside the MEG helmet was continuously monitored by means of five head position indicator (HPI) coils. A 3D digitizer (Fastrack Polhemus Inc., Colchester, VA, USA) was used to record the location of the HPI coils and the general head shape relative to three anatomical fiducials (nasion, left and right preauricular points). To track eye movements and blinks, bipolar electrodes were attached to obtain horizontal and vertical electrooculograms (HEOG and VEOG).

#### MEG preprocessing

MEG data were cleaned of external noise using the Maxfilter 2.0 software (Elekta Neuromag), applying the Signal-Space Separation (SSS) method with movement compensation (Taulu & Simola, 2006), correlation limit of 0.9 and time window of 10 seconds. Next, data were preprocessed and subsequently analysed with the FieldTrip toolbox (Oostenveld et al., 2011) running in MATLAB. Data were segmented into trial epochs from −2 to 7 s time locked to stimulus onset and then downsampled to 200 Hz. After discarding trials with muscle and jump artifacts by trialwise inspection, an Independent Component Analysis was computed. Independent components reflecting eye movements and heartbeat were identified by visual inspection of component scalp topographies, time courses and its comparison with EOG/ECG raw time-series. Noise components were removed and clean trials were visually inspected again in order to identify and remove any remaining artifact. Across participants, an average of 15% (range: 1–60 %) of all trials were discarded. The AR condition contained an average of 55 trials (range: 10-151), the IR condition contained and average of 74 trials (range: 14-135) and the CR condition contained an average of 79 trials (range: 30-109).

### QUANTIFICATION AND STATISTICAL ANALYSIS

#### Sensor space time-frequency analysis and statistics

Frequency decomposition was obtained for each trial using Fast Fourier Transform (FFT) based sliding window analysis, progressing in 50 ms steps. The window length was optimised for each frequency from 1 to 80 Hz, with a minimum of 200 ms and 5 cycles (for instance, using 500 ms/5 cycles for 10 Hz, and 200 ms/6 cycles for 30 Hz). The data in each time window were multiplied with a Hanning taper before Fourier analysis. The power values were obtained for the vertical and horizontal component of the estimated planar gradient and then combined. Finally, the resulting power maps were corrected using a baseline time window from −0.5 to 0 s, so the resulting power maps were expressed as the power relative change to this baseline period ((activity – baseline) / baseline).

To assess, in one step, which time- and frequency bins differentiate between the three memory conditions across channels, we conducted a one-way repeated measures ANOVA from −0.5 to 2s and from 1-80 Hz, using the factor Memory (CR, IR, AR). To correct for multiple comparisons across time, frequency and channels, we used a non-parametric cluster-based permutation test (Maris & Oostenveld, 2007), setting the cluster alpha at p = .05. The 10-12 Hz power time courses in Figure 2C and 2D were derived by averaging across the 10 left parietal sensors showing the maximal effect (sum of F values) within the significant cluster (Figure 2A, dashed red square). We used paired-samples T-test to statistically test the IR and the AR effects. To correct for multiple comparisons across time points, we used a non-parametric cluster based permutation test with an alpha = .05 (Maris & Oostenveld, 2007). This procedure was repeated again to check the robustness of the effects including a wider range of frequencies (8-20 Hz) (Figure S1A) and two different sensor selections: five (Figure S1B) and twenty (Figure S1C). Sensors were always selected by their maximal F value within the significant cluster.

#### Source reconstruction

To estimate the underlying brain activity for the alpha band (10-12 Hz) effects found at the sensor level, we performed source reconstruction from −.5 to 2 s. First, a regular grid of 1825 points with 10 mm spacing was created in the Colin27 MRI template (Collins et al., 1998) using Fieldtrip’s brain segmentation tools. Then, this set of points was transformed into each participant’s space using the individual headshapes derived from the 3D head digitalization. The forward model was solved with a single-shell method and the source reconstruction was performed using the linearly constrained minimum variance (lcmv) beamforming approach implemented in Fieldtrip. We constructed a common filter to ensure reliable comparison between conditions: the spatial filter’s coefficients were obtained from the average covariance matrix from all CR, IR and AR trials and then this filter was multiplied with each condition separately. To maximise the informational content of the signal (Van Veen, van Drongelen, Yuchtman, & Suzuki, 1997) while remaining within the functional definition of the alpha band, artifact-free data were initially filtered from 8 to 12 Hz with a Butterworth IIR filter as implemented in Fieldtrip. The final output consisted of a time series estimate per source location, condition and subject.

#### Source space time-frequency and statistics

Spectral analysis was performed on the reconstructed signal in the same way as in sensor space but restricted to the alpha frequency band (8-12 Hz). There are several ways to avoid a beamforming bias towards the centre of the head: although any noise bias would be the same for all conditions, we additionally baseline-corrected each condition before performing contrasts. Specifically, we computed the relative change of the power estimates using a baseline time window from −0.5 to 0 s.

To statistically test the sensor-space ANOVA effect (CR, IR, AR) in source space, we averaged source time series from 10-12 Hz and from 0.7 to 2 s (Figure 2A) and conducted a repeated-measures ANOVA (Figure 3A). To correct for multiple comparisons across source locations, we used a non-parametric cluster based permutation test with an alpha = .05.

To extract the Intersection and the Exclusive maps shown in Figure 3C and 3D respectively, we used the same source reconstructed signal (8-12 Hz) and averaged from 10-12 Hz and from 1-1.5 s. The time window choice was based on the sensor space power time courses (Figure 2C), ensuring that both the IR effect and the AR effect were present in order to avoid any bias. We then statistically tested the IR effect (IR vs CR) and the AR effect (IR vs. AR) using a paired-samples T-test. To correct for multiple comparisons across source locations, we used a non-parametric cluster based permutation test with an alpha = .05. The significant clusters obtained from these contrasts were then used to build the Intersection map and only the regions shared by both contrast (clusters) were included in the map. For visualisation purposes, the colorbar in Figure 3B shows the mean T-value of the included region (average of CR vs. IR and IR vs. AR).

For the Exclusive AR map we adopted a conservative approach. We set the alpha for the IR effect to .1 (uncorrected) and cluster corrected for the AR effect. In this way, any region showing an IR trend (even non-significant) was excluded from the AR map.

To statistically test earlier/familiarity-based IR effects (CR vs IR) in source space, we averaged source time series from 10-12 Hz and from 300 to 500 ms and conducted a paired-samples T-test (Figure S3). To correct for multiple comparisons across source locations, we used a non-parametric cluster based permutation test alpha set to .05.

To further test the IR and AR effects (Figure 3D) from 1 to 1.5s we averaged the alpha power (10-12Hz) in the regions of interest (Hippocampus and PPC) and performed a 2×3 repeated measures ANOVA with the factors Region (Hippocampus, PPC) and Memory (AR,IR,CR).

The ROI selection to extract the 10-12 Hz source power differences presented in Figure 3D and time courses presented in Figure 4 was based on the Automated Anatomical Labelling (AAL) atlas (Tzourio-Mazoyer et al., 2002) implemented in Fieldtrip. In both analyses, we averaged the sources for left and right hippocampus and PPC (including left and right inferior and superior parietal lobule) from 10 to 12 Hz. Finally, we used a paired-samples T-test to statistically test the difference between IR and AR time courses (Figure 4). To correct for multiple comparisons across time points, we used a non-parametric cluster based permutation test with an alpha = .05.

## Author contributions

Conceptualization, B.P.S. and M.W.; Formal Analysis – M.C.M.B. and B.P.S.; Investigation, B.P.S. and M.W.; Writing – Original Draft, M.C.M.B. and B.P.S.; Writing – Review & Editing, M.C.M.B., B.P.S., M.W. and R.N.H.; Funding Acquisition, B.P.S. and R.N.H.; Resources, B.P.S. and R.N.H.; Supervision, B.P.S.

## Conflict of Interest

The authors declare no competing financial interests.

## Acknowledgements

This work is supported by a Wellcome Trust/Royal Society Sir Henry Dale Fellowship (107672/Z/15/Z) and an MRC CBU intramural project grant to B.P.S.. M.C.M.B received funding by the Boehringer Ingelheim Fonds.

## Supplementary material

**Supplementary Figure 1.**
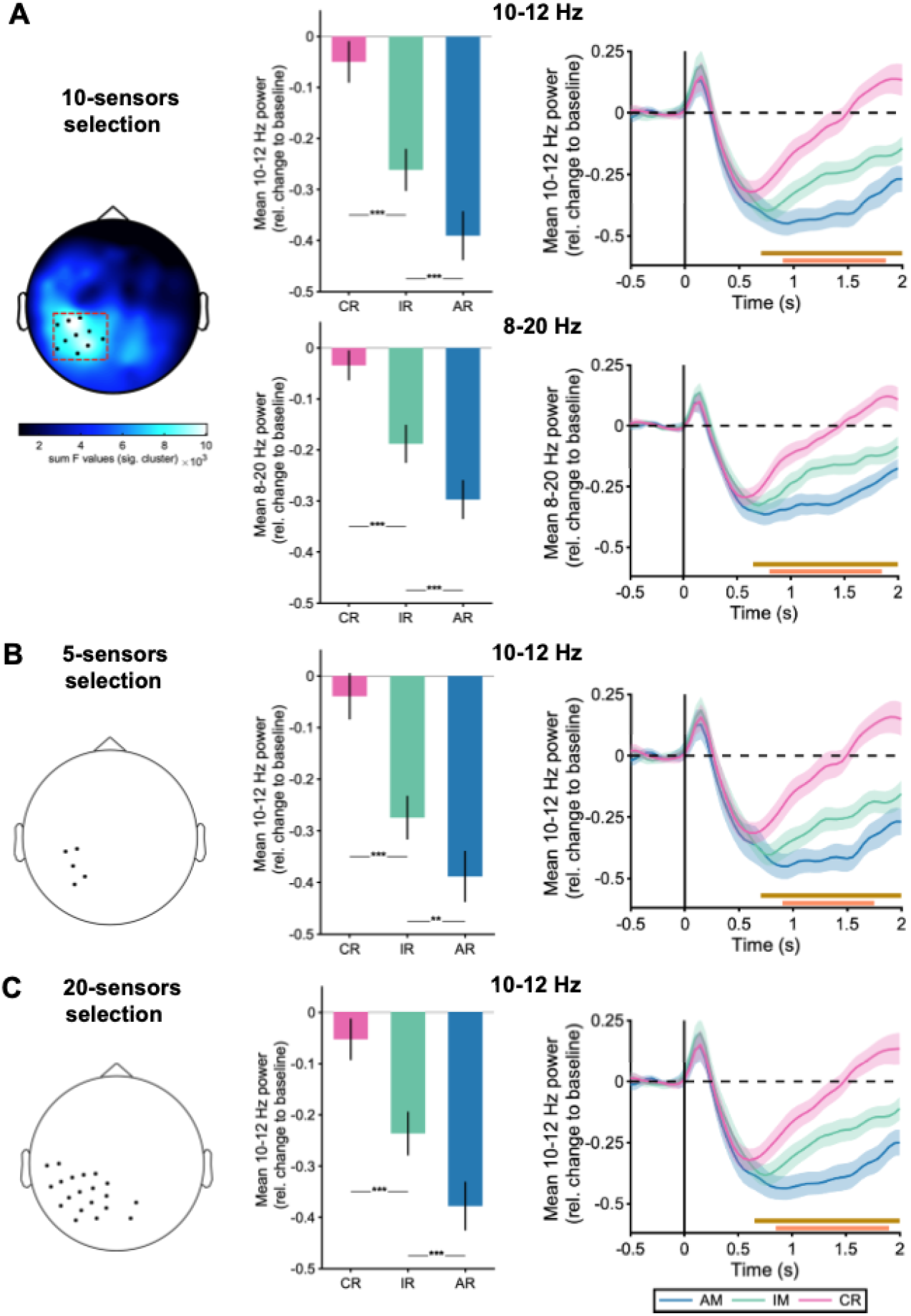
Robustness of the sensor-space results. (A) Mean (+/−SEM) power from 0.7-2 s (*left*) and stimulus-locked time courses across participants (*right*) for each memory condition in the 10-12 Hz frequency range (*top*) and 8-20 Hz (*bottom*), collapsed across the 10 sensors showing maximal F values in the main ANOVA (c.f. Figure 2A). (B) Mean (+/−SEM) alpha power from 0.7-2 s (*left*) and stimulus-locked time courses across participants (*right*) for each memory condition in the 10-12 Hz frequency range, collapsed across the 5 sensors showing maximal F values in the ANOVA. (C) Mean (+/−SEM) alpha power from 0.7-2 s (*left*) and stimulus-locked time courses across participants (*right*) for each memory condition in the 10-12 Hz frequency range, collapsed across the 20 sensors showing maximal F values in the ANOVA. ***: p < .001 and **: p < .01, paired samples t test. Brown and orange horizontal lines depict the significant clusters for item recognition memory effects (IR vs. CR) and associative recall effects (AR vs. IR), respectively (paired-samples T-tests, all p < .005).

**Supplementary Figure 2.**
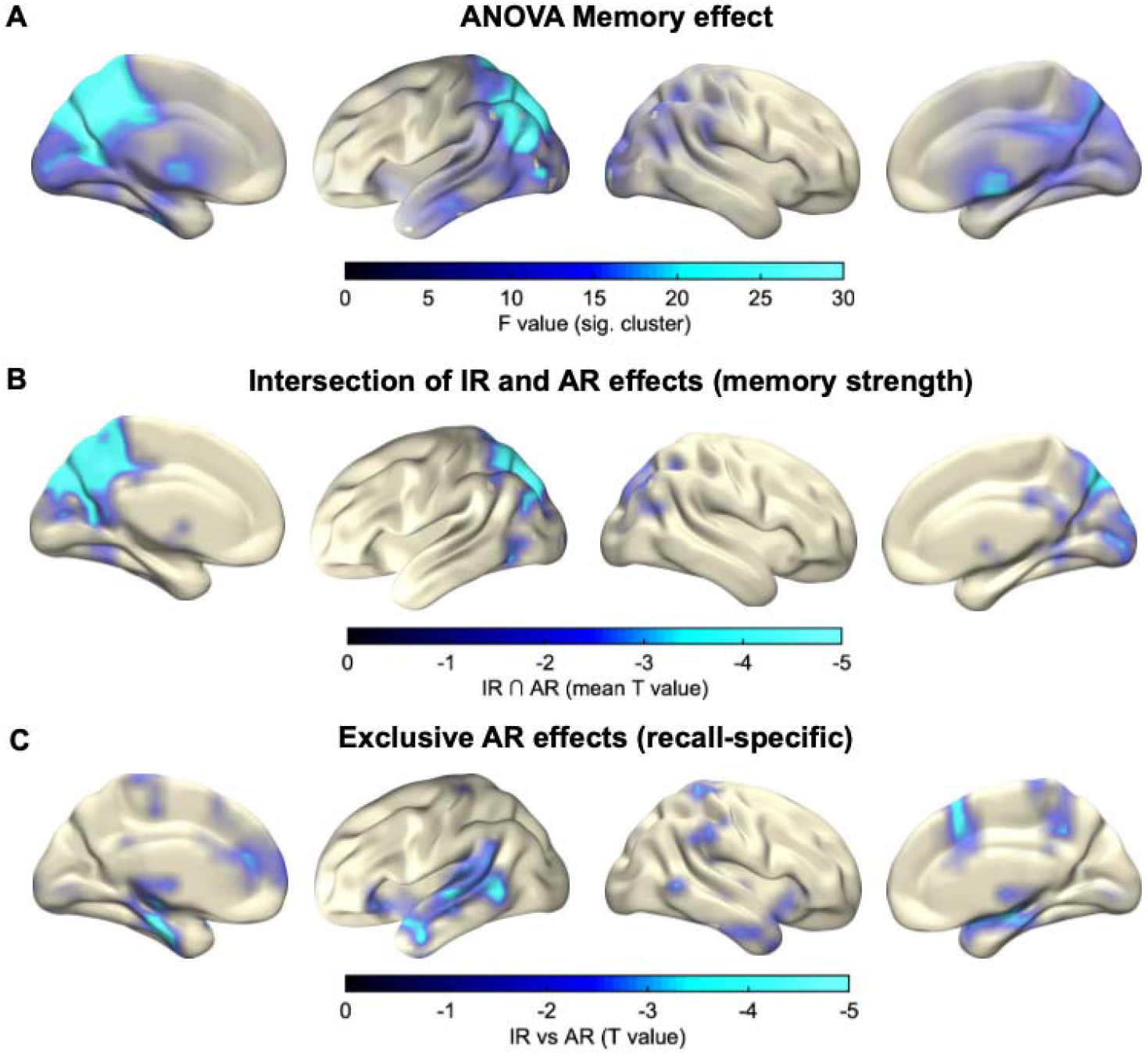
Complete view of the source reconstruction results. (A) Significant cluster resulting from the ANOVA in the 10-12 Hz alpha band from 0.5 to 2 s. (B) Regions scaling with memory strength (CR < IR < AR), revealed via inclusive masking of condition comparisons (intersection of IR vs. CR and AR vs. IR) in the 10-12 Hz alpha band and from 1 to 1.5 s. Colorbar indicates the mean T values across the IR effect (CR < IR) and the AR effect (IR < AR). (C) Exclusive AR effects (recall-specific) (1 to 1.5 s) indicates areas showing an AR effect (AR > IR, p < .05, corrected) and no IR effect (IR > CR, p < .1, uncorrected). Colorbar indicates the T value for the AR effect.

**Supplementary Figure 3.**
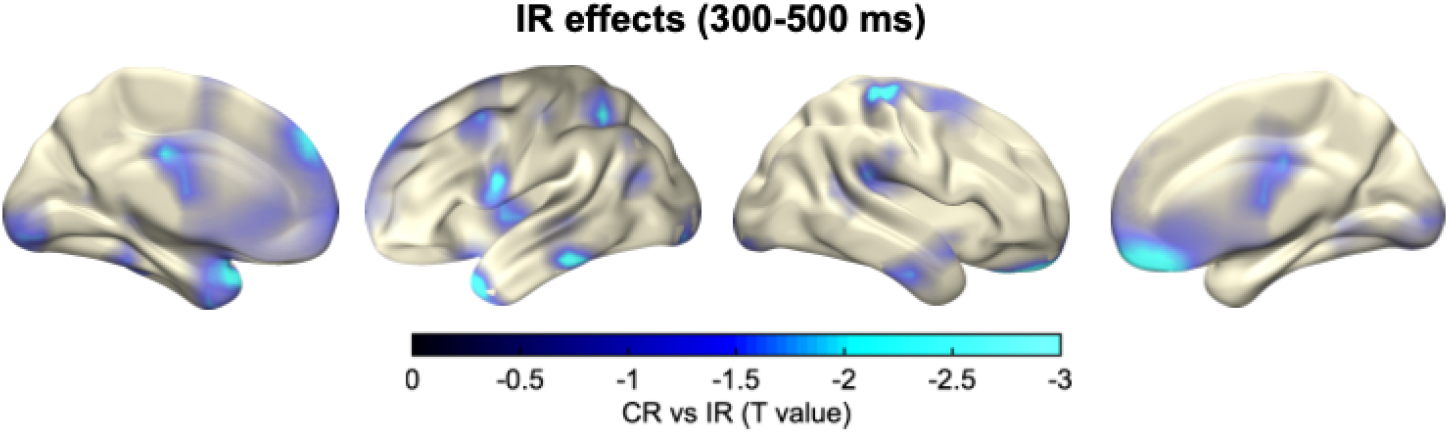
Source reconstruction of the IR effect. Significant cluster resulting from the T-test (CR vs IR) in the 10-12 Hz alpha band from 300 to 500 ms.

